# Transcriptomic Signatures of Hippocampal Subregions and Neuronal Nuclei in Active Place Avoidance Memory Maintenance

**DOI:** 10.1101/2025.09.22.677884

**Authors:** Isaac Vingan, Shwetha Phatarpekar, Victoria Sook Keng Tung, A. Iván Hernández, Oleg V. Evgrafov, Juan Marcos Alarcon

**Affiliations:** School of Graduates Studies, Program in Neural and Behavioral Sciences, State University of New York, Downstate Health Sciences University, Brooklyn, NY, USA; Institute of Genomics in Health, State University of New York, Downstate Health Sciences University, Brooklyn, NY, USA; School of Graduates Studies, Program in Molecular and Cell Biology, State University of New York, Downstate Health Sciences University, Brooklyn, NY, USA; Department of Pathology, State University of New York, Downstate Health Sciences University, Brooklyn, NY, USA; Department of Cell Biology, State University of New York, Downstate Health Sciences University, Brooklyn, NY, USA; Department of Genetics, Human Genetics Institute of New Jersey, Rutgers University, Piscataway, NJ, USA; The Robert F. Furchgott Center for Neural & Behavioral Science, State University of New York, Downstate Health Sciences University, Brooklyn, NY, USA

**Keywords:** Spatial Transcriptomics, snRNA-seq, Gene Expression, Memory, Hippocampus, Maintenance, IEG-Expressing Ensemble, Memory-associated Neuronal Ensemble

## Abstract

The gene expression changes associated with memory acquisition, consolidation and reconsolidation, all active epochs in memory formation, have been well characterized in the rodent hippocampus. Less is known, however, of the changes in gene expression supporting the maintenance of memory, particularly when it remains undisturbed or offline days after the memory experience. In this study, we used a combination of spatial transcriptomic and single nuclear RNA sequencing (snRNA-seq) to measure the gene expression changes in the dorsal hippocampus during an early phase of offline memory maintenance, 3 days after the post-training retention test of an active place avoidance memory. Through spatial transcriptomics we identified spatially regionalized differential gene expression and biological process enrichment, with CA1, CA3, and DG exhibiting differential expression of genes involved in post-synaptic function, synaptic vesicle transport, and neuronal differentiation, respectively. Notably, through snRNA-seq, differentially expressed genes detected in clusters of hippocampal neurons from the trained animal were largely defined by their down regulation of genes involved in ATP synthesis and cytoplasmic translation. With both techniques we also examined the gene expression changes in a putative subset of memory-associated neurons through the detection of eYFP mRNA in the Arc-Cre/flox-eYFP double transgenic mouse line. Amongst this population of cells, we detected a limited number of differentially expressed genes unique to each subregional population and associated with synaptic plasticity and post-synaptic signaling. Our results suggest that two overarching transcriptomic patters contribute to the functional changes in hippocampal cells during offline memory maintenance: a regional distribution of pathways linked to synaptic functions, and a reduction of metabolic activity across hippocampal sub-regions and memory-associated neuronal ensembles.

## 1 INTRODUCTION

The diversity of the transcriptomic profile across brain systems is predicted to be essential for the functioning of neuronal ensembles that support memory ^4–8^. Foundation for this prediction arises from studies that establish that the acquisition, recall and maintenance of learned information depends on the expression of synaptic plasticity mechanisms at the synapses connecting these ensembles ^9–11^; and, that changes in gene expression are an essential process to support synaptic plasticity ^7,12–15^.

In the rodent hippocampus, processing of sensory and emotive/affective contextual information has been ascribed to specific hippocampal sub-regions ^16–19^, and the formation of spatial memory is reliant on the recruitment of particular neuronal ensembles within these specific sub-regions ^20–22^. Unique neuronal operations at specific regions of the hippocampal network appear essential to the processing of memory information ^23–26^, including pattern separation and pattern completion ^27–31^, emotive and social integration, and context-content integration ^32–35^. As such, each sub-region of the hippocampal network can be implicated in neuronal computations that bear a direct functional role on specific components of memory performance ^16–18,36–39^ as well as the updating of memory information ^40–42^. Despite the enticing connection between memory-associated gene expression and memory-associated regionalization of hippocampal function, this link remains underexplored.

Our laboratory recently identified distinct transcriptomic profiles associated with the recall of an active place avoidance memory within sub-regions of the dorsal hippocampus ^43^. We found that these transcriptomic profiles were selectively expressed in cell populations identified by their expression of a single immediate-early gene (IEG; Arc, Egr1, or c-Jun), suggesting a particular transcriptomic architecture for memory recall. In the present study, we further our exploration by a characterization of memory-associated transcriptomic profiles days after the learning experience and, critically, without experimentally eliciting a memory recall.

We collected hippocampal tissue from mice trained in an active place avoidance (APA) memory task and untrained controls three days after a post-training memory retention test for RNA sequencing analyses. We test for gene expression changes through spatial transcriptomics and single nuclear RNA sequencing to compare the transcriptomes of functionally distinct hippocampal sub-regions and cells, respectively. To explore gene expression profiles in hippocampal cells associated with the active place avoidance memory experience, we used the Arc-Cre/flox-eYFP transgenic mouse ^44^ to persistently tag neurons that expressed the activity-related immediate early gene Arc during the retention test.

Our findings reveal an abundance of metabolic and energy consumption regulation processes in the hippocampus of animals trained in the active place avoidance memory task; suggesting for a differential energy balance required to support the place avoidance memory trace. A number of processes relevant for learning and memory were found in hippocampal sub-regions CA1 (e.g., postsynaptic function), CA2 (e.g., mRNA processing) and CA3 (synaptic assembly/organization) in comparison to the Dentate Gyrus (e.g., some enrichment for synaptic signaling processes).

Notably, all of the areas showed a consistent downregulation of cytoplasmic translation and metabolism. Hippocampal inhibitory neurons showed similar enrichment of biological processes than excitatory neurons, which was also observed in glial cells. Differential gene expression analysis of eYFP tagged cells identified synaptic plasticity genes in the trained cohort, but not enough to enrich biological processes; possibly due to the sparseness of the active place avoidance associated memory trace in the hippocampus.

As memory-associated neuronal ensembles dynamically reorganize throughout the phases of a memory (e.g., acquisition, reactivation, and storage) ^16–18,45,46^; our study suggest for a concomitant shift in gene expression profiles across the hippocampus to support the maintenance of a memory trace.

## 2 RESULTS

### 2.1 Animals trained in APA show robust post-training memory retention test

To investigate gene expression changes associated with offline memory maintenance in the hippocampal network, we conducted spatial transcriptomic and single nuclei RNA sequencing (snRNA-seq) studies 3 days after active place avoidance (APA) memory retention test using an Arc-Cre/flox-eYFP double transgenic mouse line (see Figure 1A for experimental timeline). Arc-expressing neurons in the brains of these Arc-Cre/flox-eYFP mice can be induced to express the fluorescent protein marker, eYFP, in a time dependent manner, following the intraperitoneal injection of 4-hydroxytamoxifen ^44^.

**Figure 1.**
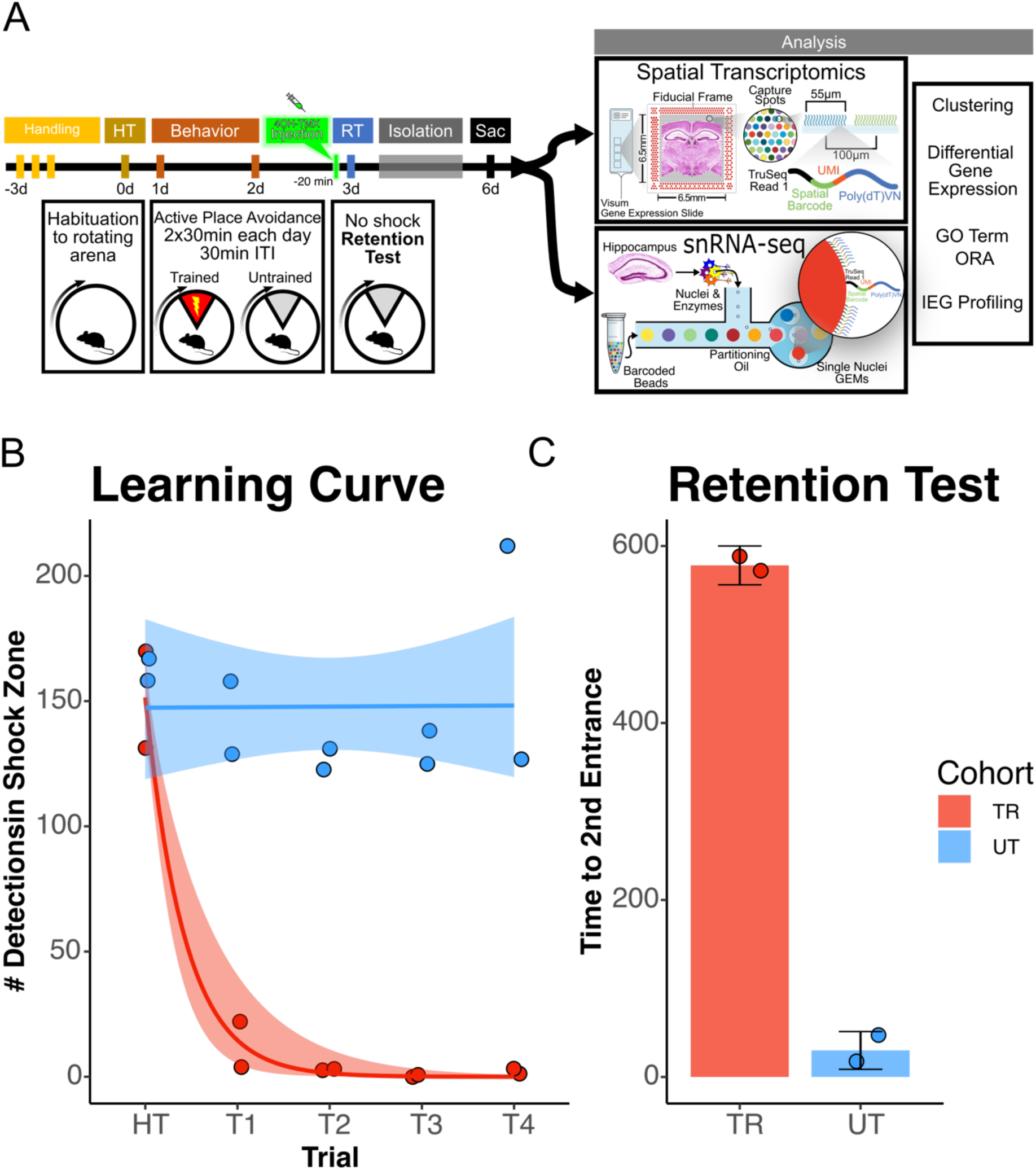
Mice trained in the APA learned to avoid the location of an unmarked shock zone. **(A)** Experimental timeline. Mice were trained in the active place avoidance paradigm over the course of 4 days following 3 days of handling. On day 0 mice were habituated to a rotating arena with no active shock zone (HT). On day 1 and 2 mice assigned to the trained behavioral condition (n =2, male) received two 30-minute trials with an active shock zone. Mice assigned to the untrained behavioral condition (n = 2, male) were not exposed to an active shock zone. On day 3, all mice received an IP injection of 4-hydroxytamoxifen to trigger permissive tagging of Arc-expressing neurons with eYFP. 30 minutes post injection, mice received a 10-minute retention test trial (exposed to the rotating arena with no active shock zone). Following the retention test, mice were returned to the colony room and single housed in a sound attenuation chamber for 3 days to limit stimulation and spurious eYFP neuronal tagging. Mice were sacrificed at end of this 3-day interval and brains were collected for further processing. **(B)** Training performance measured as number detections in the shock zone. Mice assigned to the trained behavioral condition learned to avoid the location of the shock zone indicated by the decreased number of shocked triggered. Exponential fit learning curves included with shaded error bars (SE). **(C)** Memory performance measured as time to second entrance in the retention test trial. Trained mice demonstrated higher time to second entrance during the retention test. Line and point color reflect the assigned behavioral condition (untrained = blue, trained = red).

In the active place avoidance task, mice learn to avoid a 60◦ shock zone on a rotating circular arena ^47,48^. Mice trained in the active place avoidance task with the shock zone ON (n = 2 male) exhibited a decrease in the number of detections in the shock zone during the acquisition trials (Figure 1B) and, one day later, longer re-entry times to the shock zone during the memory retention test trials (Figure 1C) relative to the mice in the untrained cohort (n = 2 male). Re-entry times to the shock zone (i.e., time to second entrance) was used as the retention test performance metric because it represents performance that is more resistant to the animal’s potential errors in self-localization once placed in the rotating arena^47–49^. After the retention test, animals were returned to their home cages and housed inside a sound attenuation chamber. Three days later, animals were taken from their home cages and immediately euthanized (5% isoflurane vaporized in 100% oxygen). Tissues harvested from the 2 trained and 2 untrained mice were utilized in both spatial transcriptomic and snRNA-seq studies.

### 2.2 Spatial transcriptomics analyses reveal regionalized patterns of gene expression 3 days after APA memory retention test

To investigate the spatial distribution of differentially expressed genes associated with active place avoidance memory maintenance across the hippocampus, we performed spatial transcriptomics on coronal sections containing the dorsal hippocampus from two trained (n = 2 male) and two untrained (n = 2 male) mice. Computational analyses of integrated capture spots in the hippocampal slices from trained and untrained animals revealed distinct clusters (seen in a Uniform Manifold Approximation and Projection (UMAP) plot) which map along anatomical boundaries in a spatial transcriptomic plot (Figure 2A, B). Spatial transcriptomic data were integrated with the Allen Brain Atlas ^8^ to provide a semi-supervised cell type annotation of each capture spot (Figure 2C, D).

**Figure 2.**
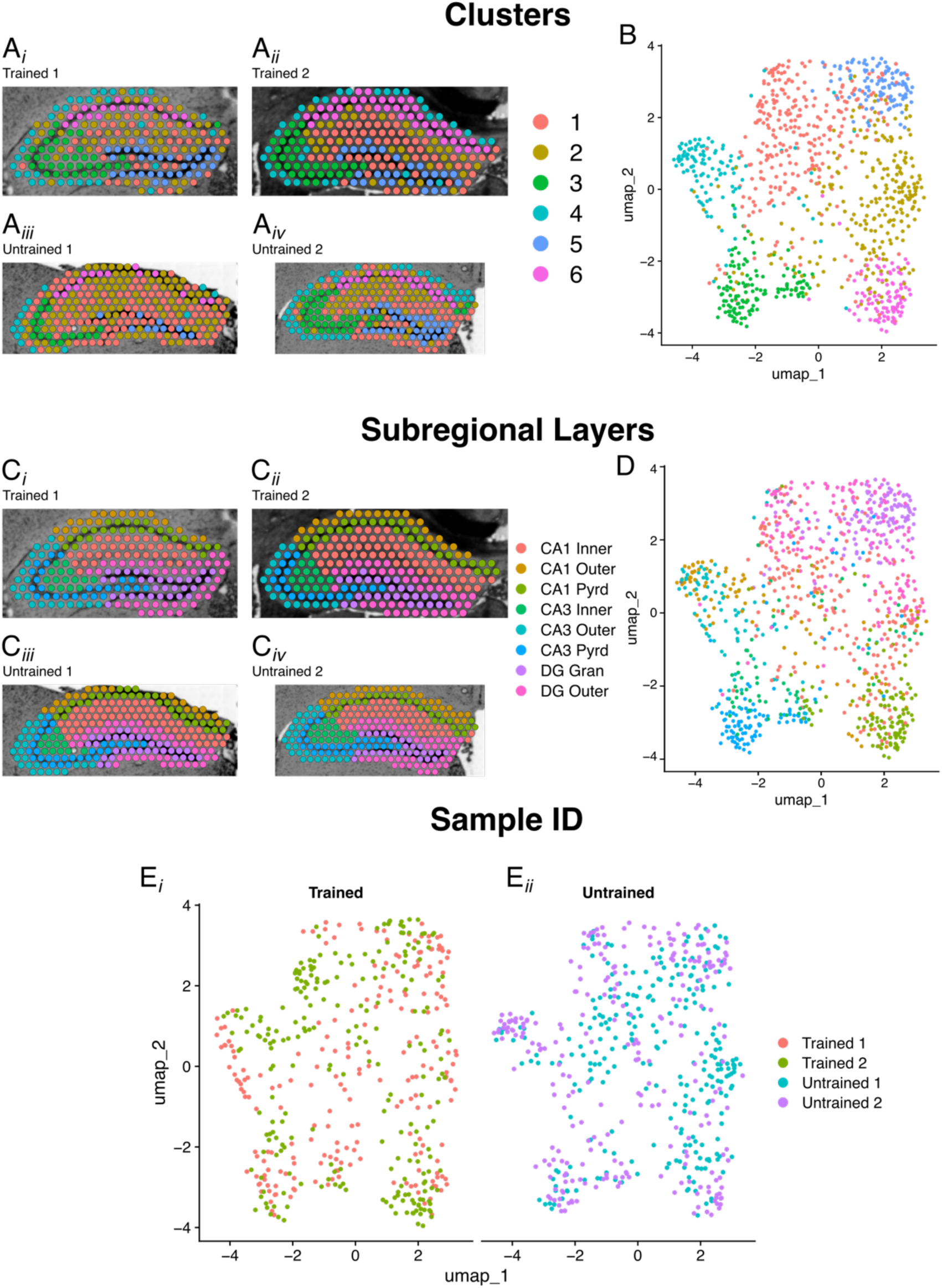
Clustering and cell-type annotation reflect anatomical boundaries of the hippocampal subregions. **(A_i_-A_iv_, B)** Spatial transcriptomics map **(A_i_-A_iv_)** and UMAP plot **(B)** of all hippocampal spots from trained and untrained mice (n = 2 per group). Discrete color scale displays unsupervised graph-based clustering of hippocampal spots. Unbiased clustering groups spots along anatomical boundaries reflecting natural transcriptomic differences between morphologically distinct regions in the hippocampus. **(C_i_-C_iv_, D)** Spatial transcriptomics map **(C_i_-C_iv_)** and UMAP plot of cell-layer annotated spatial transcriptomic maps of hippocampal spots **(D)** in samples from trained and untrained mice. Discrete color scale reflects the predominant anatomical cell layer surveyed by each capture spot which was determined through assistance by query-based cell-type annotation through data integration with the single cell RNA-seq Allen Brain Atlas cortex and hippocampus in mice. **(E_i_-E_ii_)** UMAP plots split along behavioral conditions demonstrates the relative distribution of spatial transcriptomic spots from each sample and their suitable computational integration.

Differential gene expression analysis between trained and untrained mice identified total 129 differentially expressed genes in all hippocampal spots. *Dgkz* and *Omg*, genes related to synaptic plasticity and memory, were found to be upregulated with a high fold change in this analysis^50,51^. Stratification by subregions detected 79 differentially expressed genes in the CA1, 149 in the CA3, and 76 in the DG in the trained condition. In the untrained condition, we detected 61 differentially expressed genes in the CA1, 27 in the CA3, and 47 in the DG (Figure 3A_i_-A_iii_). The genes *Plk2*, *Hs3st4*, and *Hpca*, which are related to synaptic plasticity and memory, were found to be significantly upregulated with a high fold change in the trained CA1, CA3, and DG, respectively^52–54^.

**Figure 3.**
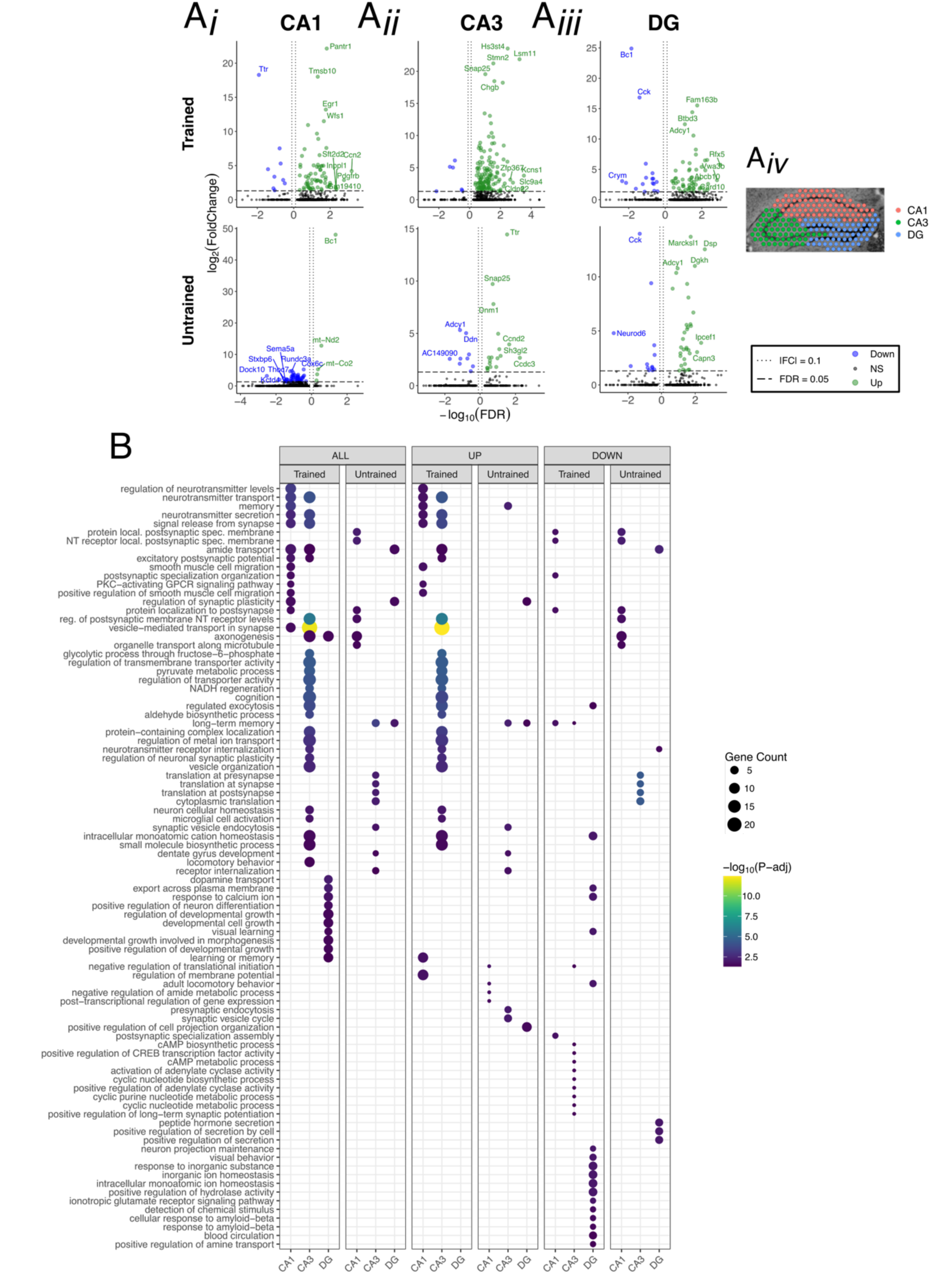
DEGs detected between trained and untrained animals within hippocampal subregions using spatial transcriptomics are enriched with genes involved in synaptic plasticity processes. **(A_i_-A_iv_)** Volcano plot of differentially expressed genes detected between combined analysis of hippocampal across trained and untrained conditions. In the trained condition, 70 differentially expressed genes were upregulated and 9 differentially expressed genes were downregulated in the CA1 **(A_i_** top panel), 143 and 6 in the CA3 **(A_ii_** top panel), and 59 and 17 in the DG **(A_iii_** top panel). In the untrained condition, 5 differentially expressed genes were upregulated and 56 differentially expressed genes were downregulated in the CA1 **(A_i_** bottom panel), 18 and 9 in the CA3 **(A_ii_** bottom panel), and 34 and 13 in the DG **(A_iii_** bottom panel). Representative spatial transcriptomic map demonstrates how capture spots were clustered in space for differential expression testing **(A_iv_**). Regional enrichment of biological processes detected amongst all differentially expressed genes (left) and stratified by up-(middle) and down-regulated (right) differentially expressed genes **(B)**. Dot color reflects the statistical significance (-log_10_(FDR)) of the biological process enrichment. Dot size reflects the number of detected differentially expressed genes mapped to the genes involved in a given biological process.

Gene ontology enrichment analyses of the differentially expressed genes detected between all hippocampal spots identified biological processes related to ATP production and ribosomal assembly upregulated in the trained condition (Figure 3B). differentially expressed genes detected in the analysis of the CA1 spots in the trained condition were enriched with genes involved in biological processes related to post synaptic function. In the CA3 of the trained condition, differentially expressed genes were enriched with genes involved in glycolysis and synaptic vesicle transport. And in the DG of the trained condition, downregulated differentially expressed genes were enriched with genes involved in neurite growth and neuronal differentiation.

### 2.3 snRNA-seq analyses reveal downregulation of genes involved in ATP synthesis and cytoplasmic translation 3 days after APA memory retention test

To complement our spatial transcriptomics data, we performed snRNA-seq on nuclei extracted from microdissected hippocampal tissue sections of trained (n = 2, male) and untrained mice (n = 2, male) to investigate the transcriptomic changes in specific populations of cells associated with the maintenance of active place avoidance memory. The tissue samples harvested in this study were microdissected from the same frozen tissue blocks used for spatial transcriptomics.

Extracted nuclei were prepared for sequencing using the 10x Chromium platform (See Methods | 7 & 8) ^55^. Sequence data from all 4 samples of nuclei were processed to remove low-quality reads, sequencing reads mapped to the genome/transcriptome and to individual beads (typically corresponding to single nuclei) to assess expression of each gene in each nucleus. Resulting data was analyzed using Seurat package. Clusters were annotated with cell types through query based annotation and integration with a subset of the single cell Allen Brain Atlas for Cortex and Hippocampus (Figure 4A,B) ^8^.

**Figure 4.**
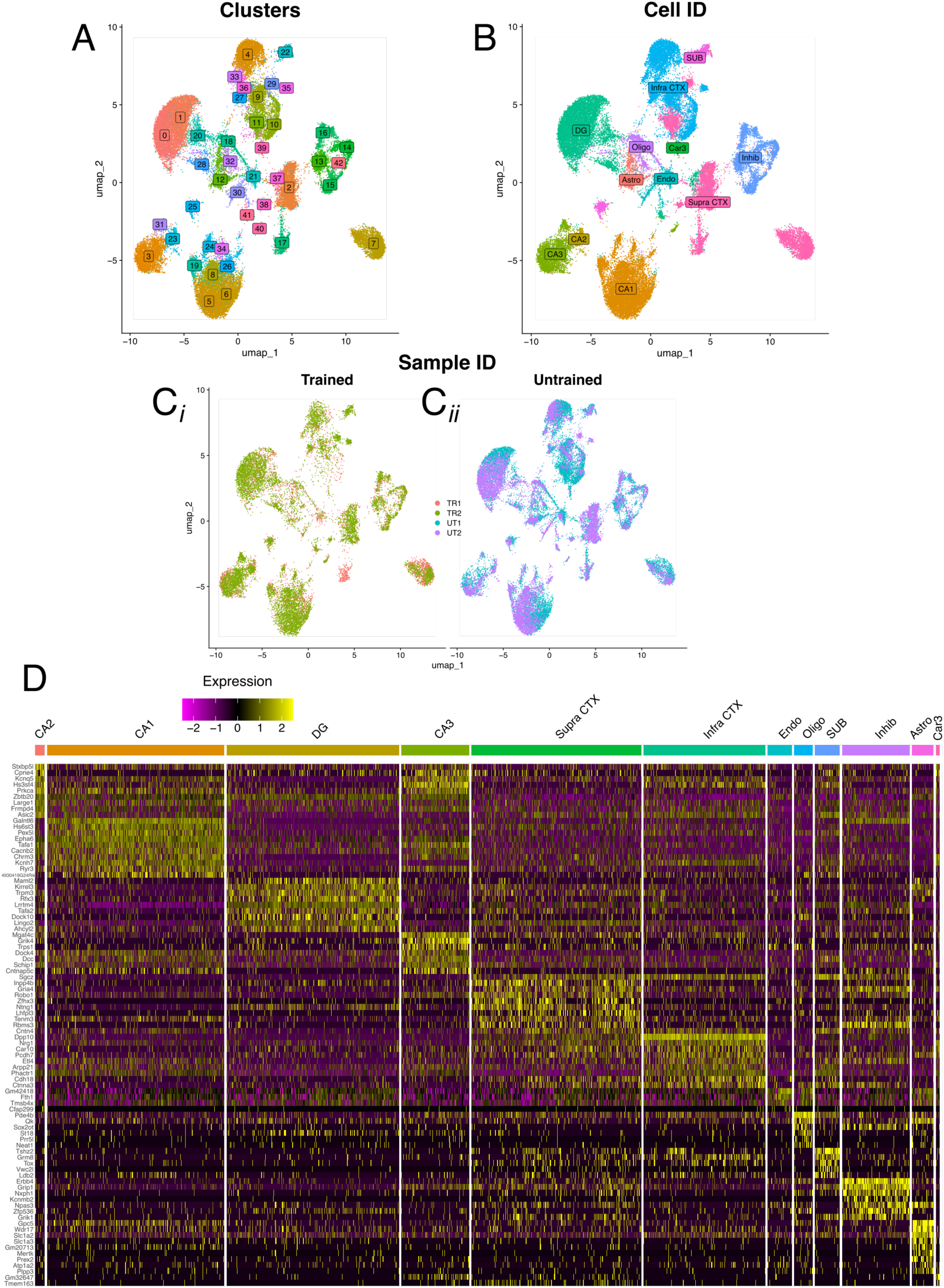
Clustering of nuclei reflects hippocampal subregional cell-types. **(A)** UMAP plot of clustered hippocampal nuclei. Discrete color scale displays unsupervised graph-based clustering (resolution = 1.5) separating nuclei into 42 distinct clusters. **(B)** UMAP plot of cell-type annotated hippocampal nuclei clusters. Discrete color scale reflects the predominant cell type surveyed by each cluster which was determined through assistance by query-based cell-type annotation through data integration with the single cell RNA-seq Allen Brain Atlas cortex and hippocampus in mice. **(C_i_, C_ii_)** UMAP plots split along behavioral conditions demonstrates the relative distribution of sequenced nuclei from each sample and their suitable computational integration. **(D)** Gene expression heatmap with each row showing the relative expression of one of the top 10 significant differentially expressed genes marking each cell-type cluster. Columns represent a random subset of 10,000 nuclei grouped by their cell type clustering and arranged by a hierarchical clustering dendrogram (not shown). Cells are colored in a continuous scale by the normalized expression of each gene in each sampled nucleus.

We analyzed a total of 45,700 sequence nuclei (16,971 from the trained animals, 28,729 from the untrained animals), comprising 12 distinct cell type clusters for: CA1, CA2, CA3, DG, Subiculum, Inhibitory neurons, Supra-granular cortical neurons, Infra-granular cortical neurons, Astrocytes, Oligodendrocytes, Endothelium and Car3 neurons. Cluster annotation was performed through query based automated annotation through integration with the Allen Brain single cell RNA-seq Atlas for Cortex and Hippocampus ^8^. Among the hippocampal nuclei we identified seven, three, and six subclusters in the CA1, CA3 and DG, respectively. Additionally, the inhibitory cell cluster contained 5 subclusters corresponding to *Sst*+, *Pvalb*+, *Vip*+, *Lamp5*+, *Sncg*+ inhibitory cell populations. Differential expression testing was performed across all cell type clusters to identify markers for each cell type. Cluster for cell types specific to the hippocampal formation were marked by *Hs6st3* in the CA1, *Stxbp5l* in the CA2, *Mgat4c*, in the CA3, *Maml2* in the DG (Figure 4D). Cells type clusters which can be found in the hippocampus and surrounding areas were marked with expression of Erbb4, in the inhibitory cell population, Slc1a2 in the astrocyte population, and Fth1 in the oligodendrocyte population.

Next, we performed differential gene expression analyses to identify the genes and molecular functions that are differentially regulated in the cells of the hippocampus. Nuclei from the CA1, CA2, CA3, DG, inhibitory, astrocyte, and oligodendrocyte clusters were each compared individually across training conditions. In total, 829 differentially expressed genes were identified in the trained-untrained comparison of CA1 nuclei, 11 in the CA2, 312 in the CA3, and 529 in the DG (Figure 5A-D). Highly significant differentially expressed genes with high fold change detected in the CA1 show differential expression of *Lrrtm1* and *Kirrel3*, which have been reported to be involved in postsynaptic development ^56,57^. Differential expression testing across training conditions in the inhibitory neuron, astrocyte and oligodendrocyte clusters detected 275, 54, and 100 differentially expressed genes, respectively (Figure 6A-C). The genes *Tenm3* and *Trpm3*, studied for their involvement in CA1 circuitry formation, were significantly upregulated with a high fold change in the inhibitory interneurons of the trained animals ^58,59^. Additionally, the gene *Ttr*, studied for its role in long term memory performance ^60–62^, was found to be significantly upregulated in every comparison of neuronal cell-type clusters across trained and untrained conditions.

**Figure 5.**
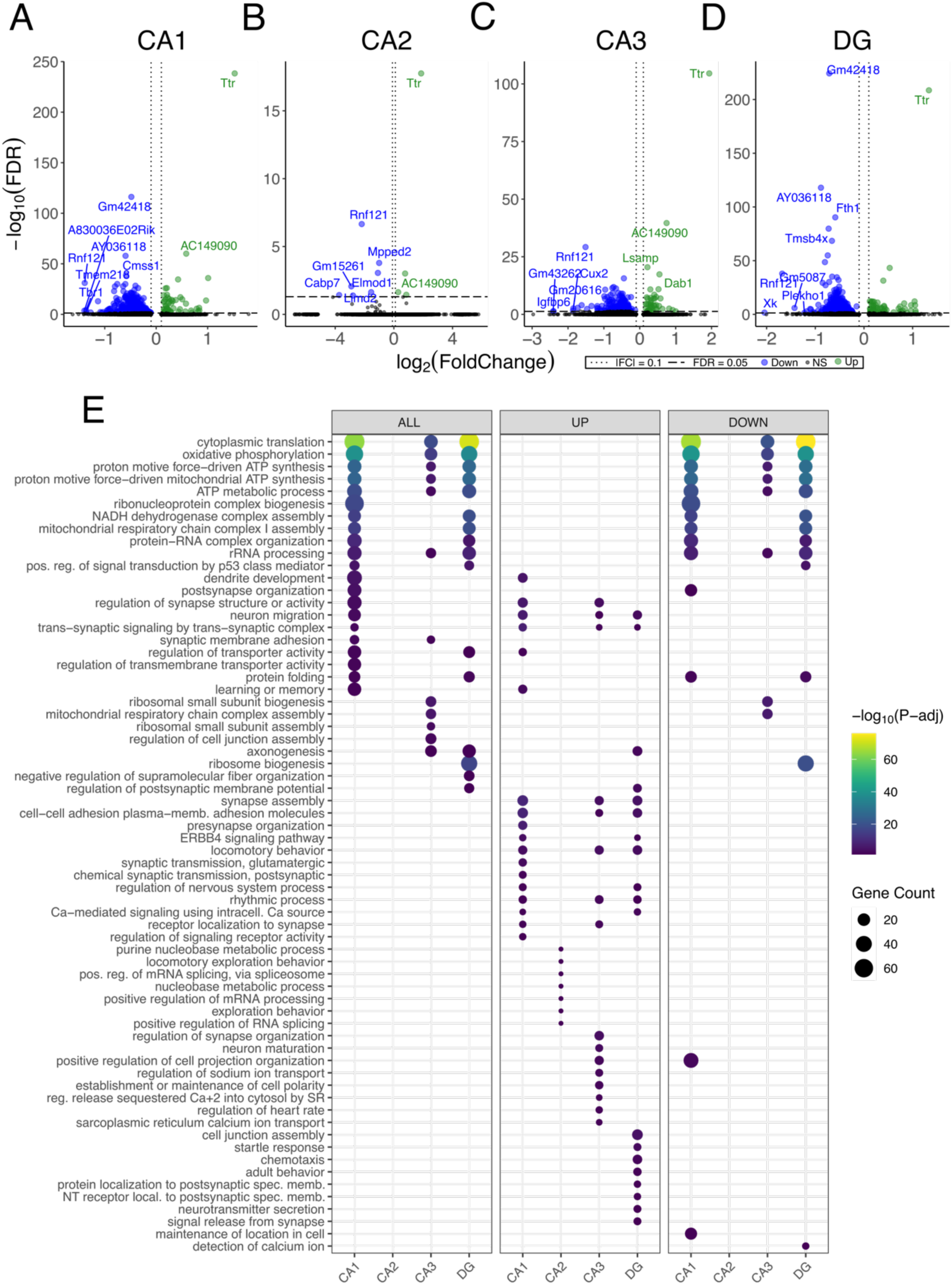
DEGs detected between trained and untrained animals within hippocampal cell-type clusters are enriched with genes involved in translation and ATP synthesis. **(A-D)** Volcano plot of differentially expressed genes between CA1 **(A)**, CA2 **(B)**, CA3 **(C)**, and DG **(D)** nuclei clusters between trained and untrained mice (n = 2 male per group). In the analysis of CA1 nuclei comparing trained and untrained conditions, 54 genes were upregulated and 775 were downregulated in the trained condition. 4 genes were upregulated and 7 were downregulated in the CA2 nuclei, 45 genes were upregulated and 267 were downregulated in the CA3 nuclei, and 65 genes were upregulated and 464 were downregulated in the DG nuclei. **(E)** Regional enrichment of biological processes detected amongst all differentially expressed genes (left) and stratified by up-(middle) and down-regulated (right) differentially expressed genes. Dot color reflects the statistical significance (-log_10_(FDR)) of the biological process enrichment. Dot size reflects the number of detected differentially expressed genes mapped to the genes involved in a given biological process.

**Figure 6.**
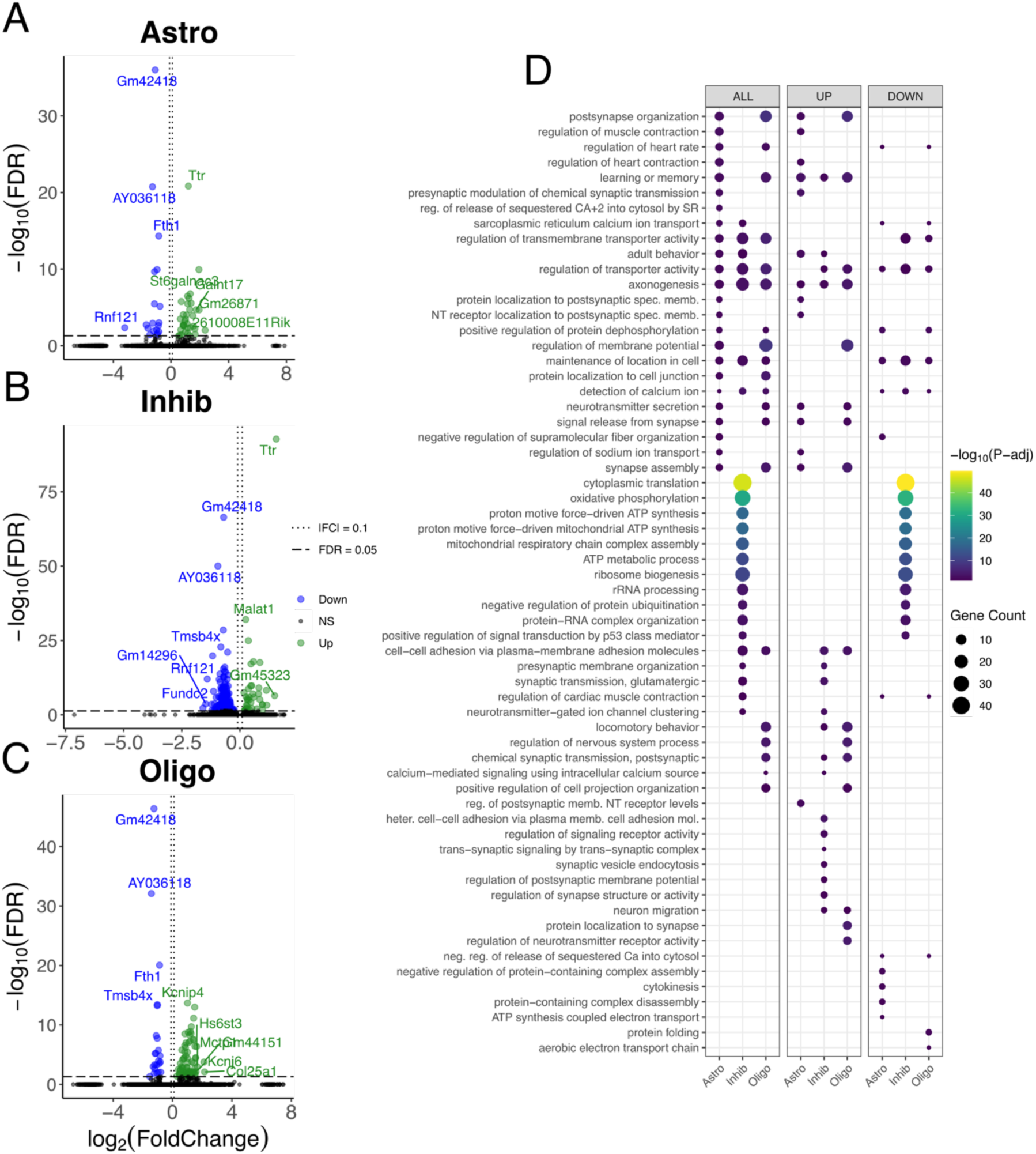
DEGs detected between trained and untrained animals within astrocyte, oligodendrocyte, and inhibitory interneuron clusters demonstrate enrichment of biological processes similar to those seen in hippocampal neuronal clusters. **(A-C)** Volcano plot of differentially expressed genes between inhibitory interneuron **(A)**, astrocyte **(B)**, and oligodendrocyte **(C)** nuclei clusters between trained and untrained mice (n = 2 male per group). In the analysis of inhibitory interneuron nuclei comparing trained and untrained conditions, 38 genes were upregulated and 237 were downregulated in the trained condition. 35 genes were upregulated and 19 were downregulated in the astrocyte nuclei, and 77 genes were upregulated and 23 were downregulated in the oligodendrocyte nuclei **(D)** Regional enrichment of biological processes detected amongst all differentially expressed genes (left) and stratified by up-(middle) and down-regulated (right) differentially expressed genes. Dot color reflects the statistical significance (-log_10_(FDR)) of the biological process enrichment. Dot size reflects the number of detected differentially expressed genes mapped to the genes involved in a given biological process.

We performed gene ontology term enrichment analyses on the lists of differentially expressed genes detected between the nuclei belonging to the trained and untrained conditions. differentially expressed genes detected in the comparison of CA1 nuclei were enriched with genes involved in biological processes related to synaptic plasticity and learning and memory (Figure 5F). Enrichment with genes related to cell-cell adhesion and ionic balances were detected in the CA3 nuclei, and synaptic signaling and cell-adhesion in the DG. We also found significant enrichment of biological processes related to mRNA processing among the upregulated differentially expressed genes in the CA2 nuclei. Notably, overrepresentation of biological processes related to cytoplasmic translation and metabolism was detected amongst the downregulated differentially expressed genes in the CA1, CA3 and DG. Similarly, biological processes related to cytoplasmic translation and ATP-production were enriched among downregulated differentially expressed genes detected in the comparison of inhibitory neurons nuclei (Figure 6D). In glial cell specific nuclei clusters, biological processes related to synaptic organization and cellular projection were overrepresented amongst the differentially expressed genes detected in both the astrocyte and oligodendrocyte nuclei clusters. Many of the biological processes significantly enriched amongst the differentially expressed genes of these glial populations corresponded to processes also found in the analyses of excitatory hippocampal nuclei.

### 2.4 Spatial transcriptomics identifies a limited number of DEGs marking eYFP-expressing spots 3 days after APA memory retention test in the trained animal

To characterize the changes in gene expression amongst putative Arc-expressing memory-associated neuronal populations in active place avoidance memory maintenance (i.e., 3 days after the active place avoidance retention test), we analyzed the eYFP-expressing population of spatial transcriptomics spots whose neurons were tagged during the active place avoidance memory retention test. Out of the 460 spots in the hippocampus from both trained animals, 79 spots (∼17%) had detectable expression of Gt(ROSA)26Sor mRNA (eYFP+). In the untrained animals, 77 eYFP+ spots out of 530 (∼14%) total hippocampal spots (Figure 7A, B). Regionally, the CA1 area contained the highest number of eYFP+ spots in the samples from both trained and untrained animals. Though, measures of spatial variability (Moran’s I) for the eYFP mRNA did not demonstrate a significant spatial distribution in either training condition.

**Figure 7.**
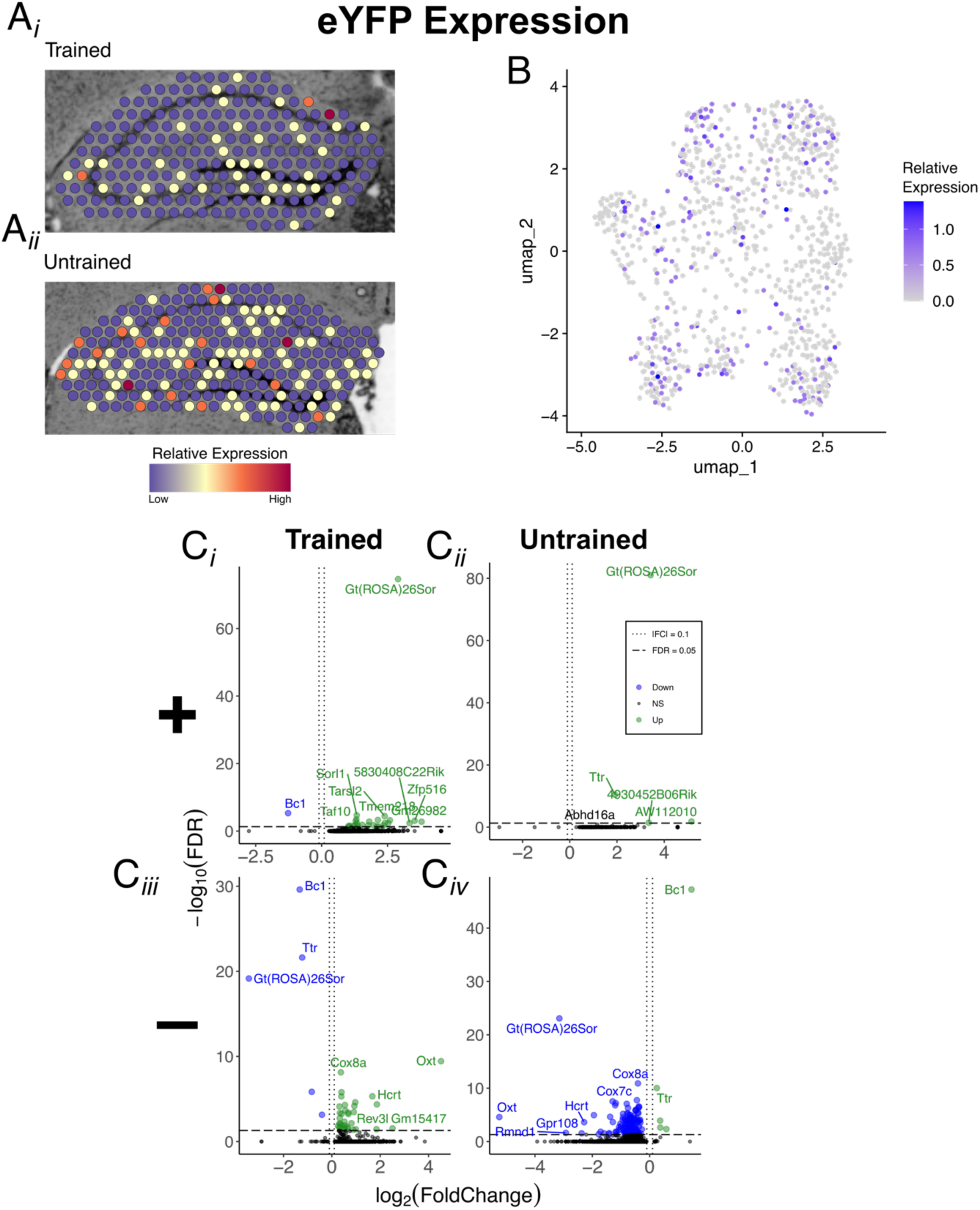
eYFP-positive spots in the trained sample are marked by genes involved in postsynaptic signal transduction. **(A_i_, A_ii_, B)** Representative spatial transcriptomic maps **(A)** UMAP plot of hippocampal spots **(B)** in the trained and untrained samples colored to reflect relative expression of the eYFP mRNA transcript, Gt(ROSA)26Sor. **(C_i_, Ciii)** Volcano plots of differential gene expression of eYFP positive spots in the trained **(C_i,_ C_iv_)** and untrained **(C_i_, Ciii)** conditions. In the analysis of eYFP*+* spots in the trained condition 26 upregulated and 1 downregulated gene were detected. In the analysis of eYFP*+* spots in the untrained condition 4 upregulated and 0 downregulated genes were detected. **(C_ii_, Civ)** Volcano plots of differential gene expression between eYFP-spots in trained **(C_ii_)** and untrained **(C_iv_)** conditions. In the analysis of eYFP*-* spots in the trained condition 35 upregulated and 5 downregulated genes were detected. In the analysis of eYFP*-* spots in the untrained condition 5 upregulated and 155 downregulated genes were detected.

Next, we performed differential expression testing to identify differentially expressed genes marking groups of hippocampal spots in each training cohort separated by eYFP positivity (Figure 7C_i_ - C_iv_). We found 27 differentially expressed genes marking the eYFP+ spots in the trained animals and 4 in the untrained animals. There were 40 and 160 marking the eYFP-spots in the trained and untrained animals, respectively. Among the differentially expressed genes marking the trained eYFP+ group of spots, a number are reported to be involved in the execution and modulation of postsynaptic neurotransmitter signals, such as *Dr1* (Dopamine receptor 1), *Rgs20* (Regulator of G-protein signaling 20), *Serpini1* (Neuroserpin) ^63–67^. Because of the low yield of differentially expressed genes marking the eYFP+ spots in both trained and untrained conditions, gene ontology term enrichment analyses did not detect significant enrichment of biological processes to report.

### 2.5 snRNA-seq detects a limited number of DEGs marking sub-regional populations of eYFP-expressing nuclei 3 days after APA memory retention test in the trained animal

To complement the spatial transcriptomics data, we next examined the eYFP-expressing population of hippocampal nuclei to assess the changes in gene expression amongst the hippocampal cells putatively associated with the active place avoidance memory. In total 785 eYFP+ nuclei were detected in this experiment, with 247 nuclei collected in the trained condition and 538 eYFP+ nuclei were detected in the untrained (Figure 8 A_i_, A_ii_). The highest proportion of eYFP+ nuclei was found in the CA1 cell type cluster for both the trained (58) and untrained (106) conditions.

**Figure 8.**
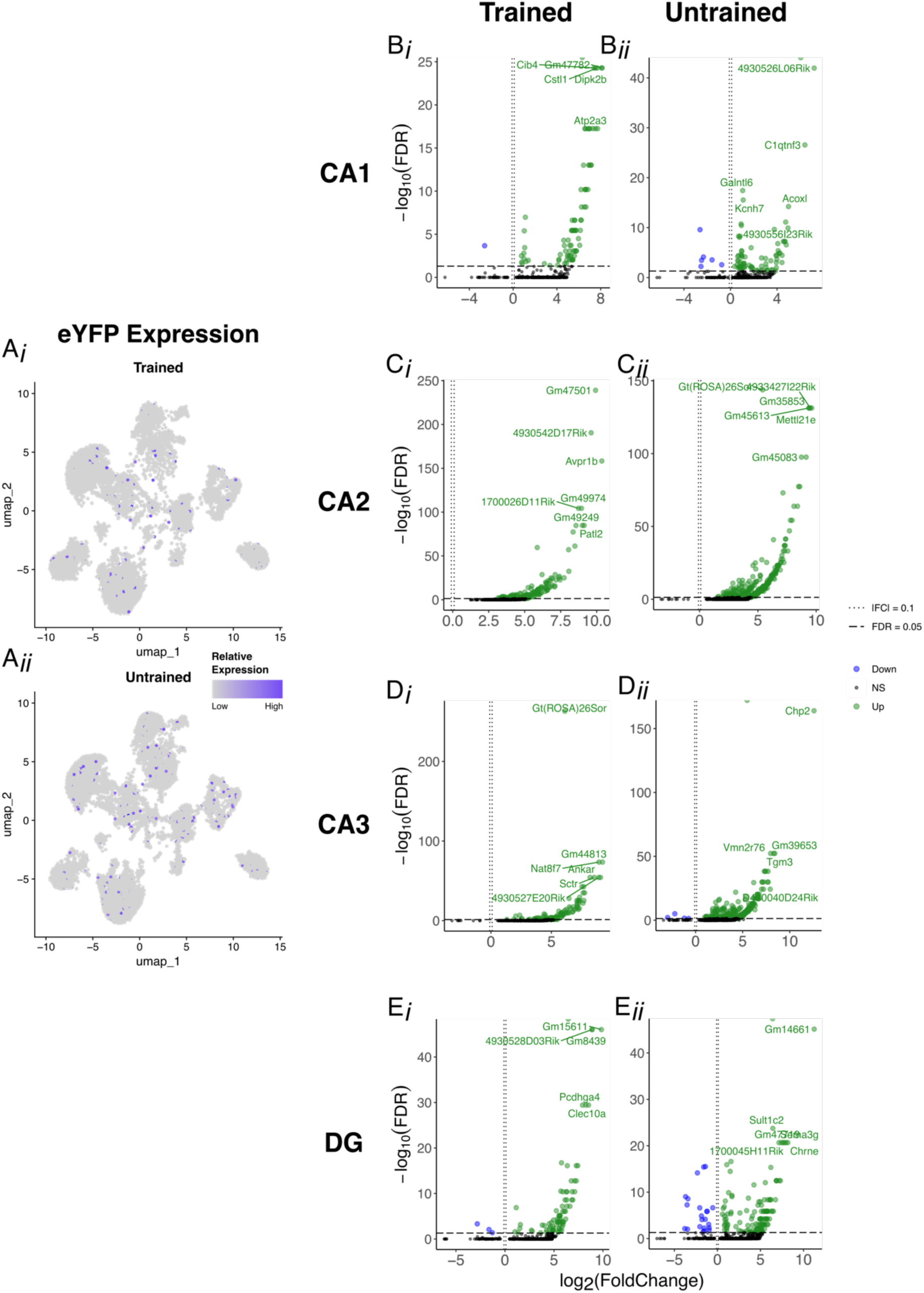
eYFP-positive nuclei in the trained sample are marked by genes involved in calcium-driven signal transduction and membrane excitability. **(A_i_, A_ii_)** UMAP plot of hippocampal nuclei split along behavioral conditions to show the distribution of eYFP+ nuclei in each cluster. Datapoints are colored with a continuous color scale to reflect the relative expression of the eYFP mRNA transcript, Gt(ROSA)26Sor. **(B_i_ – E_ii_)** Volcano plots of differential gene expression marking eYFP+ nuclei in the trained condition for CA1 nuclei **(B_i_)**, CA2 nuclei **(C_i_)**, CA3 nuclei **(D_i_)**, and DG nuclei **(E_i_). (B_ii_ – E_ii_)** Volcano plots of differential gene expression marking eYFP+ nuclei in the untrained condition for CA1 nuclei **(B_ii_)**, CA2 nuclei **(C_ii_)**, CA3 nuclei **(D_ii_)**, and DG nuclei **(E_ii_).**

Differential expression testing was performed first comparing all eYFP+ samples collected from the trained and untrained animals. In this analysis, 7 differentially expressed genes were found to mark the eYFP+ cells in the sample from trained animals, and 14 in the sample from untrained animals. The analysis was further stratified by region, comparing all eYFP+ nuclei amongst nuclei clusters that principally belonged to the hippocampal formation (CA1, CA2, CA3, DG).

We detected 83, 161, 105, and 83 differentially expressed genes marking the eYFP+ nuclei in the CA1, CA2, CA3, DG of the trained animals, respectively (Figure B_i_-E_i_). In the untrained animals, we detected 83, 325, 261, and 161 differentially expressed genes marking the CA1, CA2, CA3, DG, respectively (Figure B_ii_-E_ii_). Among the differentially expressed genes marking the CA1 eYFP+ nuclei in the trained animals, we detected significant differential expression of the genes *Slc8a1* (Solute Carrier Family 8 Member A1), *Ppp2r2b* (Protein Phosphatase 2 Regulatory Subunit B Beta), and *Prckb* (Protein Kinase C Beta) all of which are associated with calcium driven signal transduction in the neuron ^68–72^. Genes involved in neuronal membrane excitability, such as *Grik4* (Glutamate Ionotropic Receptor Kainate Type Subunit 4) and *Kcnq5* (Kv7.5), were found amongst the differentially expressed genes marking the eYFP+ nuclei from the trained animal in the CA3 cluster. In the same manner as seen for the spatial transcriptomics study, gene ontology term enrichment analyses did not detect significant enrichment of biological processes amongst the differentially expressed genes marking the regionalized subpopulations of eYFP+ nuclei.

## 3 DISCUSSION

In this study, we used a combination of spatial transcriptomics and snRNA-seq to identify changes in gene expression within major subregions of the dorsal hippocampus 3 days following a post-training active place avoidance memory retention test. We leveraged an Arc-Cre/flox-eYFP double transgenic mouse line to enhance our investigation of the gene expression changes supporting memory maintenance by also directly probing the neurons tagged at the time of the active place avoidance memory retention test.

### 3.1 Technical and Methodological considerations

Both spatial transcriptomics and snRNA-sequencing experiments detected a regionally distinct enrichment of biological process canonically ascribed to neuronal function with memory. Yet, the concordance between these processes in each region was less than expected across the two analyses. This could be accounted for by the methodological differences inherent to each experiment. (1) Spatial transcriptomics probes a single 10µm thick tissue section containing the dorsal hippocampus. Because tissue samples for snRNA-seq were collected from the same brain after collecting the spatial transcriptomic slices, they originated from a more posterior hippocampal region. Anatomical differences alone might be sufficient to explain the molecular differences observed between spatial transcriptomics and snRNA-seq analyses, as indicated by the distribution of gene expression profiles native to the mouse hippocampus along the dorsal-ventral axis ^5^. (2) There are molecular differences in how the two transcriptomic methodologies sample mRNA. Spatial transcriptomics, with its utilization of 55µm diameter capture spots, is well equip to sample intact tissue microenvironments without single cell specificity ^73,74^. This is contrasted by snRNA-seq’s sampling of mRNA in and around single nuclei while removing cytoplasmic mRNA that was transported to the soma or neuronal projections at an earlier time point ^75–77^. With this in mind, it is possible that with spatial transcriptomics we are surveying the net effect of all cells and their projections within a given microenvironment, while with snRNA-seq we are observing the molecular changes spatially and temporally located around the cell nucleus at the time of collection.

### 3.2 Memory maintenance is supported by subregional gene expression profiles and metabolic dampening in the hippocampus

The functions neurons perform during offline memory maintenance are poorly understood ^78^. Much of the literature investigating the molecular mechanisms of memory put major emphasis on the immediate-recall and late-remote phases of memory, often with the addition of behavioral probe that will reactivate the consolidated memory trace ^78–86^. MAPK and PKA signaling pathways have been highlighted as a critical molecular pathway in the hippocampus during the retrieval phase of memory ^80^. In contrast, studies of the molecular mechanisms in the hippocampus after a recall-inducing experience and a period of sleep highlights the role of the regulation of trans-synaptic signaling in the DG ^87^. The data in the present study complement these recent reports by showing that the molecular landscape in memory associated brain systems in the days following memory recall is diverse and dynamic. In this memory epoch, neuronal systems are transitioning from active consolidation and reconsolidation – canonically defined by waves of new protein synthesis – to a role in offline memory maintenance and distribution of the hippocampal memory trace across cortical brain regions ^82,84,88–91^. The evidence in this study indicates that, though the hippocampal subregions of the trained animal are still involved in processes that modulated synaptic efficacy and membrane excitability, by and large, hippocampal neurons are defaulting to conservative state that is catabolically and energetically depressed.

The subregions of the hippocampus participate in different features of memory performance as the memory trace is consolidated and maintained amongst their cells ^33,92–98^. We found this evidence that these regionalized roles in memory maintenance influence the transcriptomes of the cell in each of the major hippocampal subregions. The field has long known about the importance of the organization of the post-synaptic terminal in memory and cognitive function^99^. This notion is supported by the detection of biological process enrichment related to post-synaptic membrane specialization in the CA1 of the trained animal in our spatial transcriptomic study. Synaptic vesicle transport has also been associated with memory consolidation, which we also find enriched among the differentially expressed genes of the CA3 of the trained animal in the spatial transcriptomic study ^100,101^. Single nuclei RNA-seq revealed biological processes related to ATP synthesis and translation enriched in the downregulated markers for each neuronal cell-type cluster in the hippocampus (including inhibitory neurons and glial cells). This finding was unexpected. It did not involve canonical synaptic plasticity processes or follow a regionally distinct transcriptome pattern. But, it may indicate an unexplored role of energy efficiency and reduced protein synthesis during offline memory maintenance in the absence of an experimentally-induced memory trace reactivation ^102,103^.

### 3.3 Select few molecular features define the memory associated eYFP expressing neuronal ensembles in memory maintenance

We leverage the persistent fluorescent tagging of neurons in the Arc-Cre/eYFP double transgenic mouse model to investigate the transcriptomic changes in a putative subset of the memory-associated neuronal ensemble. Following the injection of 4-hydroxytamoxifen, neurons that were sufficiently activated to express the immediate early gene Arc during the retention test would be induced to persistently express eYFP mRNA and protein, which could be detected at a later time point. This animal model, and others like it, have been widely utilized to investigate the cells strongly activated during learning or memory related epoch – the memory-associated neuronal ensembles ^44,104–108^. To date, only a handful of studies have focused on elucidating the molecular profiles defining this critical population of neurons ^7,13,15,109–114^.

In the present study we identified the genes marking populations of eYFP expressing neurons. Through spatial transcriptomics we found upregulation of *Dr1* and *Gad2*, pointing toward the role of dopamine signaling and inhibition supporting the memory trace in the microenvironment surrounding the memory-associated neuronal ensemble ^64,115^. Through snRNA-seq, we identified distinct lists of differentially expressed genes marking the hippocampal subregionalized eYFP positive neuronal populations in the trained mouse, plausibly indicating that eYFP positive neurons express different genes depending on their anatomical location to maintain memory-related information at the molecular level. Though no biological processes were enriched among these differentially expressed genes, we did detect up regulation of the gene *Prkcb* in the CA1 which has been show to enhance synaptic efficacy in memory, and *Grik4* in the CA3 which has been shown to regulate synaptic inhibition in memory ^72,116^.

Interestingly, the limited number of genes marking and shared between the eYFP population did differ from those identified in previous transcriptomic investigations of memory-associated tagged neurons ^7,112,113^. On one hand, and as discussed previously, spatial transcriptomics is well equipped to profile tissue microenvironments, such as the collection of cells that contain at least one eYFP expressing neuron, and snRNA-seq profiles mRNA in and around the nucleus, likely not detecting the reportedly critical dendritic mRNAs. On the other, the sparsity of differential gene expression marking these populations of cells could be a reflection of the sparsity of the memory-associated neuronal ensemble itself ^45,117,118^. Notably, studies using immediate early gene expression as proxy measurements of memory-associated neuronal ensembles explore the gene expression properties of these cells after the learning or consolidation phases of memory ^7,15,109,111–114^. There is evidence to support that the memory-associated neuronal ensemble reduces in size with time since memory acquisition ^44,119,120^. Could it be possible that the sparse differentially expressed genes that mark eYFP tagged tissue spots and nuclei are evidence of this engrammatic sparsing that begins as a newly consolidated memory progresses in its transition to a more offline/remote state?

### 3.4 Memory maintenance is an understudied and molecularly complex phase of memory

We utilized spatial transcriptomics and snRNA-seq in an integrative investigation of the hippocampal transcriptional architecture associated with offline memory maintenance 3 days following the post-training active place avoidance retention test. Regional patterns of gene expression and biological process enrichment were detected in both spatial and nuclear transcriptomic techniques. With spatial transcriptomic the CA1, CA3 and DG hippocampal subregions were defined by post-synaptic function, synaptic vesicle transport, and neuronal differentiation respectively. With snRNA-seq we found that all neuronal nuclei from the hippocampus were defined by their downregulation genes belonging to biological processes involved in ATP synthesis and cytoplasmic translation. Through the detection of eYFP mRNA, which is persistently tagging a putative subset of the memory associated neuronal ensemble, we identified a sparse collection of differentially expressed genes marking these cells.

The present work explores an understudied epoch of memory, offline maintenance. Classically, the phase of memory preceding this epoch, consolidation, is defined by increases in long term potentiation and new protein synthesis ^121–125^. The phase of memory temporally downstream of offline maintenance is remote storage, which is through to involve a systems consolidation of the memory trace through its transfer out of the hippocampus and into cortical brain networks ^82,126^. The fact that our data deviate from the classical predications of memory-associated cellular changes might be an early reflection of the transition of the memory trace out of the hippocampus. Additionally, the hippocampal memory trace is theorized to function like an index, with few specialized set of neurons directing network activity toward the brain regions (e.g., PFC, retrosplenial cortex) containing relevant traces of a given memory experience ^127,128^; ^129–131^. The sparse population of eYFP tagged neurons could be evidence of the contracting memory trace into this residual index, one that is transcriptionally depressed yet primed for reactivation should the appropriate network activity arise.

Another important consideration in relation to the molecular nature of the memory trace pertains to the nature of the memory experience. Performance in the active place avoidance paradigm depends on numerous factors that inform active place avoidance memory, such as cue based navigation, behavioral inhibition, cognitive control, and aversive conditioning ^132–136^. The execution of these processes by a neuronal network, like the hippocampus, is likely to induce a diverse set of demands on the activated neurons. This robust activity would set in motion waves of transcription and translation that establish quasi-permanent changes in the synaptic and global properties of these neurons. While the molecular consequences of this activity may be easily detected immediately following activation/reactivation, the molecular impression of the memory trace alone at a later time point may be much more difficult to discriminate from the background basal neuronal activity.

## METHODS

### 1 ANIMALS

A total of 18 adult ArcCreERT2::eYFPflx mice ^44^ with C57BL/6 genetic background aged 3-4 months were utilized across all experimental cohorts. Bulk RNA sequencing experiments utilized 3 male and 3 female mice, 2 females and 1 male were assigned to the trained behavioral condition, and 1 female and 2 males were assigned to the untrained behavioral condition (see 2.2 Behavior for details). Spatial transcriptomics experiments for the study of memory 1 hour after reactivation utilized 2 male mice, 1 mouse was assigned to each behavioral condition. Spatial transcriptomics and snRNA-seq experiments for the study of memory 3 days after reactivation utilized the same 4 male mice for both experiments, 2 were assigned to each behavioral condition. And RT-qPCR experiment utilized 22 male mice, with 12 mice assigned to the trained behavioral condition and 10 assigned to the untrained. Prior to experimental onset mice were bred in-house at the SUNY Downstate Health Sciences University vivarium (Brooklyn, NY, USA). Mice were housed in groups of two to five per cage. Beginning on the first day of training and testing, mice were single housed in shoebox cages in a sound attenuation cubicle (Med Associates). For every experimental run, a minimum of two mice —each undergoing the same behavioral conditioning— were simultaneously housed in the sound attenuation cubicle. Ad libitum food and water was provided. Mice were randomly assigned to behavioral cohorts before the start of the experiment. Mice were handled daily for 3 days prior to the start of the experiment to reduce anxiety and improve voluntary approach. All animal procedures proposed are approved by, and will be performed following, the Institutional Animal Care and Use Committee guidelines at SUNY Downstate Health Sciences University.

### 2 BEHAVIOR

All procedures were performed in compliance with the Institutional Animal Care and Use Committee of the State University of New York, Downstate Health Sciences University. ArcCreERT2::eYFPflx male mice were trained in a hippocampus-dependent two-frame active place avoidance task. The place avoidance system consisted of a 40-cm diameter arena with a parallel rod floor that could rotate at 1 rpm. The position of the animal was tracked using PC-based software (Tracker, Bio-Signal Group Corp., Brooklyn, NY) that analyzed 30-Hz digital video images from an overhead camera. Mice in the trained condition learned the “Room+Arena−” task variant. Place avoidance of a 60° zone was reinforced by a constant current foot shock (60 Hz, 500ms, 0.2mA) that was scrambled (5-poles) across pairs of the floor rods. Rotation of the arena would carry the mouse into the shock zone unless the animal actively avoided the zone. Entering the shock zone for more than 500 ms triggered shock. Additional shocks occurred every 0.5 seconds until the animal left the shock zone. Measures of place avoidance were computed by TrackAnalysis software (Bio-Signal Group Corp., Brooklyn, NY). The number of detections of the animal in the shock zone was tracked as the performance metric for the training trials.

On day one, mice assigned to the trained behavioral condition (trained mice) received a 30-min trial with the shock off to habituate to the rotating arena. Across the next two days the animals experienced four training trials, with two 30-minute trials a day with the activated shock with a 40 min inter-trial-interval. Control (untrained mice) experienced identical training conditions, except for the shock always being off. The number of detections of the animal in the shock zone was tracked as the training performance metric for the training trials. Memory retention performance was assessed 24 hours after the final training session in a 10-minute retention test with the shock off. Latency to the animal’s second entrance into the shock zone was tracked as the memory retention performance metric for the retention test. Previous research indicates the latency to second entrance as a valid behavioral metric for spontaneous memory recall and a the retention test performance metric because it represents performance that is more resistant to the animal’s potential errors in self-localization once placed in the rotating arena ^47–49^.

Mice assigned to the study investigating memory 1 hour after reactivation were taken from their home cage 60 minutes after active place avoidance retention test and immediately euthanized (5% isoflurane vaporized in 100% oxygen). Mice assigned to the study investigating memory 3 days after reactivation received an intraperitoneal injection of 4-hydroxytamoxifen under anesthesia (5% isoflurane vaporized in 100% oxygen) 20 minutes before the start of the retention test trial. After the retention test, animals were returned to their home cages and housed inside a sound attenuation chamber. Three days later, animals were taken from their home cages and euthanized.

### 3 IN VIVO ARC-DRIVEN NEURONAL TAGGING

We use the Arc-Cre/flox-eYFP double transgenic mouse to identify neurons that undergo strong neuronal activity 3 days following memory reactivation in the retention test trial. This strong neuronal activity drives *Arc* promoter-mediated, Cre recombinase-regulated expression of eYFP ^44^. The first transgene is a construct that contains an Arc gene promoter that mediates the synthesis of a Tamoxifen-sensitive Cre-Recombinase. The second transgene contains a floxxed R26R bacterial promoter that mediates the constitutively active expression of eYFP. In the presence of exogenous 4-hydroxytamoxifen, the Cre-Recombinase recombines the R26R promoter into an active configuration, thus triggering the expression of eYFP in neurons that undergo Arc promoter-mediated gene expression (Figure M2). This permits the identification of Arc-driven (eYFP+) neuronal ensembles in the brain in a temporally regulated manner. Intraperitoneal injection of 4-hydroxytamoxifen (1.5 mg/ml) makes eYFP expression permissible within 20 minutes. After injection, mice rest in their home-cage for 20 minutes prior to the retest trial. After the trial, mice are returned to their home-cage for 20 minutes, and then are returned to a single-housed sound attenuation chamber for 3 days to avoid spurious activity while eYFP expression is still permissible.

**Figure M.2:**
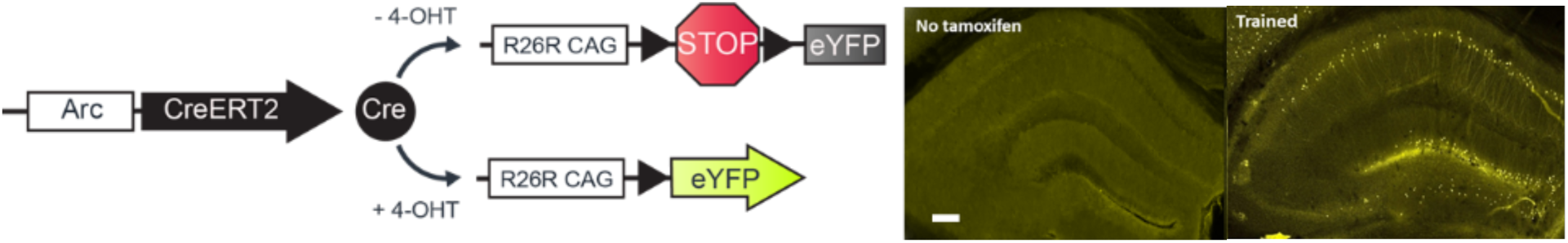
Double Transgenic Arc-Driven eYFP Expression System. Left: Arc-Cre/flox-eYFP double transgene and Cre recombinase activation by 4-hydroxytamoxifen. right: No eYFP signal is detected in mice that were not injected with 4-hydroxytamoxifen (image is from an active place avoidance trained mouse). Right: Only after 4-hydroxytamoxifen injection, we observe eYFP labeling of cells (please see fig. 2). Bar: 500 mm.

### 4 TISSUE PREPARATION FOR SPATIAL TRANSCRIPTOMICS

60 minutes after retention test, mice were euthanized and brains were extracted and immediately prepared for snap freezing in cuvettes of Tissue-Tek Optimal Cutting Temperature compound (Sakura Finetek USA, Torrance, CA, United States) floating in a bath of methyl-butane chilled by liquid nitrogen. Brain blocks were cryosectioned at −20°C ^141^ and 10 µm thick coronal sections containing the dorsal portion of the hippocampus were obtained (bregma −1.7 and −2.2 mm ^142^). Collected tissue sections are trimmed to fit within the 5 mm x 5 mm capture area on the Visium gene expression slide. Sections were mounted on a prechilled Visium gene expression slide or a Tissue Optimization slide.

For Optimization, all tissue sections were collected from a single tissue. For Visium gene expression, slide capture areas contained one section per biological sample. Slides were fixed in prechilled methanol before staining using the Visium spatial tissue optimization protocol ^143^ or the Visium spatial gene expression protocol ^144^.

Tissues permeabilization time on the gene expression slide was set to 18 minutes based on our tissue optimization results. Images were taken according to the Visium Spatial Gene Expression Imaging Guidelines ^145^. Brightfield histology images for the Visium Tissue Optimization and Gene Expression slides were taken using the Leica Aperio CS2 Slide Scanner at 20x magnification. Tissue optimization fluorescent images were taken on a Zeiss LSM 800 (555nm LED,75% intensity, and 200ms exposure).

mRNA was extracted and libraries were prepared following the Visium Spatial Gene Expression User Guide ^144^. They were loaded at 300 pM and sequenced on a NovaSeq 6000 System (Illumina) using a NovaSeq S4 Reagent Kit (200 cycles, catalog no. 20027466, Illumina), targeting a sequencing depth of approximately 2.0 × 10^8^ read-pairs per sample. Sequencing depth was determined through a calculation of a suggestion 50,000 read pairs per spatial capture spot^144^. Our tissue samples covered roughly 80% of the 5000 spatial capture spots within the Visium fiducial frame (0.8 × 5000) × 50,000 = 2.0 × 10^8^. Sequencing was performed using the following read protocol: read 1: 28 cycles; i7 index read: 10 cycles; i5 index read: 10 cycles; and read 2: 91 cycles.

### 5 SPATIAL TRANSCRIPTOMICS PREPROCESSING

Raw RNA sequencing data obtained from NovaSeq 6000 were systemically processed to ensure high-quality outputs for downstream analyses. Raw sequencing data in BCL format were converted using *bcl2fastq2* v2.20 to FASTQ format, and simultaneously demultiplexing to assign sequences to their respective samples based on both i5 and i7 index sequences. Following the conversion, these FASTQ files were subsequently processed with *TrimGalore* v.0.6.5 to automate quality, adapter trimming and perform quality control.

For the alignment and the quantification of gene expression, *Space Ranger* v2.0.0 (10x Genomics) was used to generate and quantify raw gene-barcode matrices from RNA sequencing data. Alignment and quantification of Visium spatial gene expression data utilizes the same tools implemented in the analysis of single cell RNA-seq data sets. Data were aligned to refdata-gex-mm10-2020-A, a mouse genome index provided by 10x Genomics, which is an annotated version of mm10 genome assembly. The gene-barcode matrices identified the number of unique molecular identifiers (UMIs) associated with each gene for each cellular barcode, allowing the quantification of transcript abundance.

After the alignment and quantification, Space Ranger toolset was utilized to normalize the gene expression data within the raw gene-barcode matrices to account for any technical variations. Finally, we also utilized Space Ranger to align gene expression data with spatial coordinates derived from histological images of each sample, facilitating the visualization of transcriptional activity to specific tissue location.

### 6 SPATIAL TRANSCRIPTOMICS ANALYSIS

Gene expression outputs from Space Ranger were analyzed using the software package Seurat v5 in the R coding environment ^146,147^. Spatial gene expression data were cropped to only include the capture spots within the dorsal hippocampus. Using default Seurat commands, data from each sample were normalized, transformed (using SCTransform v2) and integrated. The integrated data set were further normalized and clustered (resolution = 0.8) in principal component analysis space. Clustered data can be visualized through UMAP projections of the capture spots or in the 2D space of the tissue section. Cell type annotation was performed by integrating our data with a random subset of 10,000 hippocampal cells from the Allen Brain Atlas Cortex and Hippocampus Single Cell Taxonomy ^8^. Predicted cell-type identities were calculated and optimized based on gene expression similarity and anatomical location.

Capture spots were grouped along two different criteria to calculate differential gene expression in our samples. First, capture spots belonging to the whole hippocampus and each anatomical cell layer in the DG, CA3 and CA1 regions were compared across training conditions. Next, within each training condition, hippocampal capture spots were separated into two groups based on the detectable (>0) expression of the immediate early gene (e.g. Arc+ and Arc-groups). immediate early gene expressing and not expressing groups of spots were compared both within training conditions and across training conditions. Differential gene expression was calculated using the Wilcoxon rank sum test (the default method for Seurat). Genes with an of FDR below 0.05 were considered differentially expressed genes/transcripts. gene ontology term enrichment and overlap analyses on detected differentially expressed genes were performed as described above for bulk RNA sequencing differentially expressed genes.

Expression data for 23 immediate early genes from all hippocampal spots evaluated through pairwise Pearson’s Correlations using the R-language base stats package. Pearson’s correlation coefficients between all immediate early gene pairs were calculated within samples, alongside P values to assess significance. Heatmaps of the correlation matrix were plotted using the R-language package, ComplexHeatmap.

### 7 NUCLEI ISOLATION AND COUNTING FOR SNRNA-SEQ

Nuclei are extracted from minced frozen hippocampi to form clean single nuclei suspensions using a variation of the “Daughter of Frankenstein Protocol”, a 10x genomic community protocol developed by the Martelotto Lab (Martelotto, L., Daughter of Frankenstein protocol for nuclei isolation from fresh and frozen tissues using OptiPrep® continuous gradient V.2. Protocols.io, 2021.). Extracellular matrices are ruptured by douncing minced fresh frozen hippocampal sections on ice in a dilute buffered solution of IGEPAL-40 detergent until homogenous. The homogenate is filtered through a 70µm mesh cell strainer into a clean tube and spun down gently (500xg) in a refrigerated microcentrifuge.

The nuclei pellet is purified through density gradient centrifugation. The pellet is then resuspended in a 1mL of G25 solution (25% iodixanol) and underlaid under concentrated G25 solution to form a density gradient in a microcentrifuge tube to gene. Tubes are spun a 4°C. The pellet is resuspended in the wash buffer and strained through a 40µm mesh cell strainer into a fresh microcentrifuge tube.

The nuclei are counted manually in a Neubauer Hemocytometer. The nuclei suspension is stained with trypan blue dye to visualize nuclei under optical microscopy. Nuclei concentration was approximated from hemocytometry counts using the following equation: (Total Nuclei Counted)/(#of Squares Counted)×Dilution Factor × 10,000. Counted nuclei suspensions were diluted to an appropriate concentration where roughly 8,000 nuclei were contained in the loading volume of 75 µL.

### 8 DROPLET-BASED NUCLEI SEPARATION

Nuclei were prepared for sequencing using the Chromium kit and protocol CG000204 ^55^. Nuclei from each sample were loaded into the chromium chip, along with partitioning oil, and barcoded gel emulsion beads. Microfluidic channels pair nuclei with barcoded gem beads in a droplet of oil filled with reverse transcription master mix. The resultant suspension is incubated through a series of reverse transcription and library preparation steps according to the manufacture’s specifications. Cleaned libraries were loaded at 300 pM and sequenced on a NovaSeq 6000 System (Illumina) using a NovaSeq S4 Reagent Kit (200 cycles, catalog no. 20027466, Illumina), targeting a sequencing depth of approximately 250–400 10 × 106 read-pairs per sample. Sequencing depth was determined through a calculation of a suggested 20,000 read pairs per cell ^55^. Sequencing was performed using the following read protocol: read 1: 28 cycles; i7 index read: 10 cycles; i5 index read: 10 cycles; and read 2: 91 cycles.

### 9 SINGLE NUCLEI RNA SEQ PROCESSING

Raw RNA sequencing data obtained from NovaSeq 6000 were systemically processed to ensure high-quality outputs for downstream analyses. Raw sequencing data in BCL format were converted using bcl2fastq2 v2.20 to FASTQ format, and simultaneously demultiplexing to assign sequences to their respective samples based on both i5 and i7 index sequences. Following the conversion, these FASTQ files were subsequently processed with TrimGalore v.0.6.5 to automate quality, adapter trimming and perform quality control.

For the alignment and the quantification of gene expression, Cell Ranger v8.0 (10x Genomics) was used to generate and quantify raw gene-barcode matrices from RNA sequencing data. Alignment and quantification of Chromium expression data utilizes the same tools implemented in the analysis of single cell RNA-seq data sets. Data were aligned to refdata-gex-mm10-2020-A, a mouse genome index provided by 10x Genomics, which is an annotated version of mm10 genome assembly. The gene-barcode matrices identified the number of unique molecular identifiers (UMIs) associated with each gene for each cellular barcode, allowing the quantification of transcript abundance. After the alignment and quantification, Cell Ranger toolset was utilized to normalize the gene expression data within the raw gene-barcode matrices to account for any technical variations.

### 10 snRNA-SEQ ANALYSIS

Gene expression outputs from Cell Ranger were analyzed using the software package Seurat v5 in the R coding environment ^146,147^. Using default Seurat commands, data from each sample were individual cleaned and evaluated for quality control. Sequenced barcodes with fewer than 500 genes per barcode were filtered out to confounding debris-filled droplets, along with barcodes detecting greater than 5% mitochondrial RNA transcripts. Doublets were removed statistically using the R language package DoubletFinder ^148^. Individually cleaned sample data sets were normalized, transformed (using SCTransform v2) and integrated along behavioral conditions. The integrated data set were further normalized using SCTransform v2 and graph-based clusters (resolution = 1.5) were calculated in principal component analysis space. Cell type annotation was performed through query-based automated annotation by integrating our data with a random subset of 50,000 hippocampal and cortical cells from the Allen Brain Atlas Cortex and Hippocampus Single Cell Taxonomy ^8^. Predicted cell-type identities were calculated and used to optimize the cell-type annotation of graph-based clusters.

Nuclei were grouped and compared in two different ways to conduct differential expression testing in our samples. Clusters of nuclei with the same cell-type identity were compared pairwise across behavioral groups using the Seuart command FindMarkers(). Nuclei were also grouped based on the detectable expression (>0) of the eYFP mRNA Gt(ROSA)26Sor. eYFP+ and eYFP-nuclei belonging to hippocampal cell type clusters (CA1, CA2, CA3, DG, Inhibitory, Oligo, Astro) were compared for differential testing across behavioral groups using the Seurat command FindAllMarkers(). Differential gene expression was calculated using the Wilcoxon rank sum test (the default method for Seurat). Genes with an of FDR below 0.05 were considered differentially expressed genes/transcripts. Gene ontology term enrichment and overlap analyses on detected differentially expressed genes were performed as described above for bulk RNA sequencing differentially expressed genes.

